# An integrated approach to study chromatin and transcriptional dynamics throughout the cell cycle without pharmacological inhibition

**DOI:** 10.1101/2022.06.18.496661

**Authors:** Daria Shlyueva, Sarah Teed-Singer, Yingqian Zhan, Richard Koche, Kristian Helin

## Abstract

The cellular and nuclear states are constantly changing and closely linked to DNA replication and cell division, defining cell cycle phases. Pharmacological inhibition is a widely used strategy to enrich for specific phases to study them in isolation. However, chemical-induced cell cycle synchronization may influence chromatin openness and gene expression compared to unperturbed cell cycle states. Here, we use different synchronization methods to obtain G1, S and G2/M phases from mouse embryonic stem cells (mESCs) and profile their chromatin accessibility and post-translational modifications as well as steady and nascent transcripts levels. We demonstrate that synchronization with thymidine and nocodazole leads to dramatic changes in chromatin compaction and gene expression, which only partially resemble *bona fide* cell cycle-dependent differences. Blocking with CDK1 inhibitor results in efficient synchronization and evokes moderate changes in chromatin accessibility, resembling G2/M phase. Correlative analysis suggests that cell cycle-specific variations in chromatin openness in unperturbed mESCs are mediated by CDK1 substrates and CTCF. This integrated strategy can be applied to most cultured cell lines under any conditions and holds the potential to further our understanding of cell cycle-dependent susceptibility to external cues and chromatin regulation.

## INTRODUCTION

A cell can be described by a set of properties such as DNA content, chromatin profiles, transcription, proteomic and metabolic states that define specific cellular states, termed cell cycle phases. The timely progression from one state to another is essential for cell division, growth, development, and this progression is severely deregulated in malignant cells (Murray and Kirschner, 1989; Otto and Sicinski, 2017). Despite dramatic changes in cellular and nuclear states during DNA replication and mitosis, the unstimulated cells faithfully reestablish chromatin and transcriptional properties after each round of cell division. (Abramo et al., 2019; Behera et al., 2019; Festuccia et al., 2019; Petryk et al., 2018; Stewart-Morgan et al., 2019; Teves et al., 2016). At the same time, external signals can be differently read and interpreted by cells in distinct phases of the cell cycle leading to different outcomes of cell fate (L. Liu et al., 2017; 2019; Pauklin and Vallier, 2013). To link cell cycle, transcriptional regulation, and potential cell fate decisions in various conditions and genetic backgrounds, we need a comprehensive approach that allows isolation of any cell cycle phase and integration with major genomics techniques.

Pharmacological inhibition of cell cycle progression is one of the most used approaches to capture cells at a specific phase as defined by DNA content. The inhibitors typically target S phase by affecting DNA replication or G2-to-M transition via varying mechanisms. More specifically, nocodazole binds to tubulin and inhibits microtubule assembly, arresting cells in M phase (De Brabander et al., 1976). A small-molecule inhibitor of CDK1 blocks its kinase activity and prevents transitioning into M phase (Vassilev et al., 2006). Treatment with thymidine leads to an imbalance in the dNTP pool and, subsequently, to a block of DNA replication and synchronization at the G1/S boundary (Bjursell and Reichard, 1973). There is a limited number of reports that characterize the effects of the inhibitors on cell functions and genetic stability (Cooper et al., 2006; Vassilev et al., 2006; Yiangou et al., 2019) as well as the influence of cell cycle arrest on the proteome (Ly et al., 2015; 2017). However, we have not been able to identify published studies that compared transcriptional and chromatin states in chemically induced cell cycle phases to the corresponding phases in asynchronous cells.

An alternative to drug-induced synchronization is the isolation of specific cell cycle phases by sorting using individual cell cycle reporters (Festuccia et al., 2019; Friman et al., 2019), FUCCI (fluorescent <u>u</u>biquitination-based cell cycle <u>in</u>dicator) cell lines (Sakaue-Sawano et al., 2008; Singh et al., 2013) or by sorting fixed cells based on DNA content. The FUCCI system requires the establishment of a cell line with oscillating fluorescent reporters and allows to distinguish between G1, S and G2/M phases (Pauklin and Vallier, 2013; Sakaue-Sawano et al., 2008; Singh et al., 2013). Such a method is limited to a specific cell line and does not foster discrimination of quiescent cells or stain for additional nuclear factors (e.g., to separate M phase). The most adaptable approach is the sorting of fixed cells based on DNA content. It can be applied to any cultured cell line and be combined with pharmacological inhibition and staining for intracellular molecules. To get more precise discrimination of S phase from other phases, cells can be labeled with 5-ethynyl-2′-deoxyuridine (EdU), a thymidine analog. EdU, incorporated into a newly synthesized DNA strand, can be detected by click reaction with a fluorochrome-conjugated azide (Darzynkiewicz et al., 2011). Click reagents are small, not immunoreactive, and do not require harsh cellular permeabilization. The labeling is widely used to study DNA replication (Macheret and Halazonetis, 2019) and to profile native chromatin on nascent and parental DNA strands (Petryk et al., 2018; Stewart-Morgan et al., 2019). While it is well-established that EdU labeling does not affect chromatin structure and transcription, it is not clear whether the sorting of fixed and EdU-labeled cells is compatible with major functional genomics assays.

Here, we characterize *bona fide* cell cycle changes in chromatin accessibility and gene expression and address how they compare to changes in pharmacologically synchronized cells. We optimize and integrate cell cycle profiles, based on EdU labeling, in mouse embryonic stem cells (mESCs) with chromatin openness (ATAC-seq) and steady state mRNA (3’ mRNA-seq) for unperturbed cell cycle phases (G1, S and G2/M). We identify genomic loci with differential accessibility during the cell cycle and find that a subset of these is predicted to bind CDK1 substrates and CTCF. Using cell cycle-specific chromatin and transcriptional states, we show that treatment with nocodazole and CDK1 inhibitor, but not thymidine, leads to partial synchronization of chromatin accessibility and not transcription. Finally, we demonstrate that cell cycle-based sorting is compatible with the profiling of histone modifications using CUT&Tag (Kaya-Okur et al., 2019) as well as metabolic RNA sequencing, SLAM-seq (Herzog et al., 2017).

## RESULTS

### Chromatin accessibility and gene expression are dynamic throughout the cell cycle

To profile the cell cycle, we employed the labeling of replicating DNA with 5-ethynyl-2′-deoxyuridine (EdU) in mESCs. We tested the compatibility of EdU and DAPI stainings with ethanol (EtOH) or formaldehyde (FA) fixation and did not observe any discrepancies in FACS profiles (Figure 1A). Ethanol led to better preservation of RNA as assessed by 28S to 18S rRNA peaks ratio (Figure S1A). However, FA fixation was able to maintain chromatin structure similar to non-fixed cells as seen from the size patterns of the ATAC-seq libraries (Figure S1B). FA-fixed ATAC-seq data also had a similar signal-to-noise level as accessibility data from non-fixed mESCs (Figure S1C, S1D), however, with more short reads (around 200 bp) (Figure S1E). Short pulses of EdU (10 min) have been used previously to study molecular events associated with nascent DNA strand: restoration of transcription and chromatin accessibility (Stewart-Morgan et al., 2019), nucleosomal occupancy (Gutiérrez et al., 2019) and chromatin maturation (Kliszczak et al., 2011). Therefore, labeling followed by click reaction does not appear to affect transcription and chromatin structure. This suggests that the choice of fixation should be mainly determined by downstream applications.

**Figure 1.**
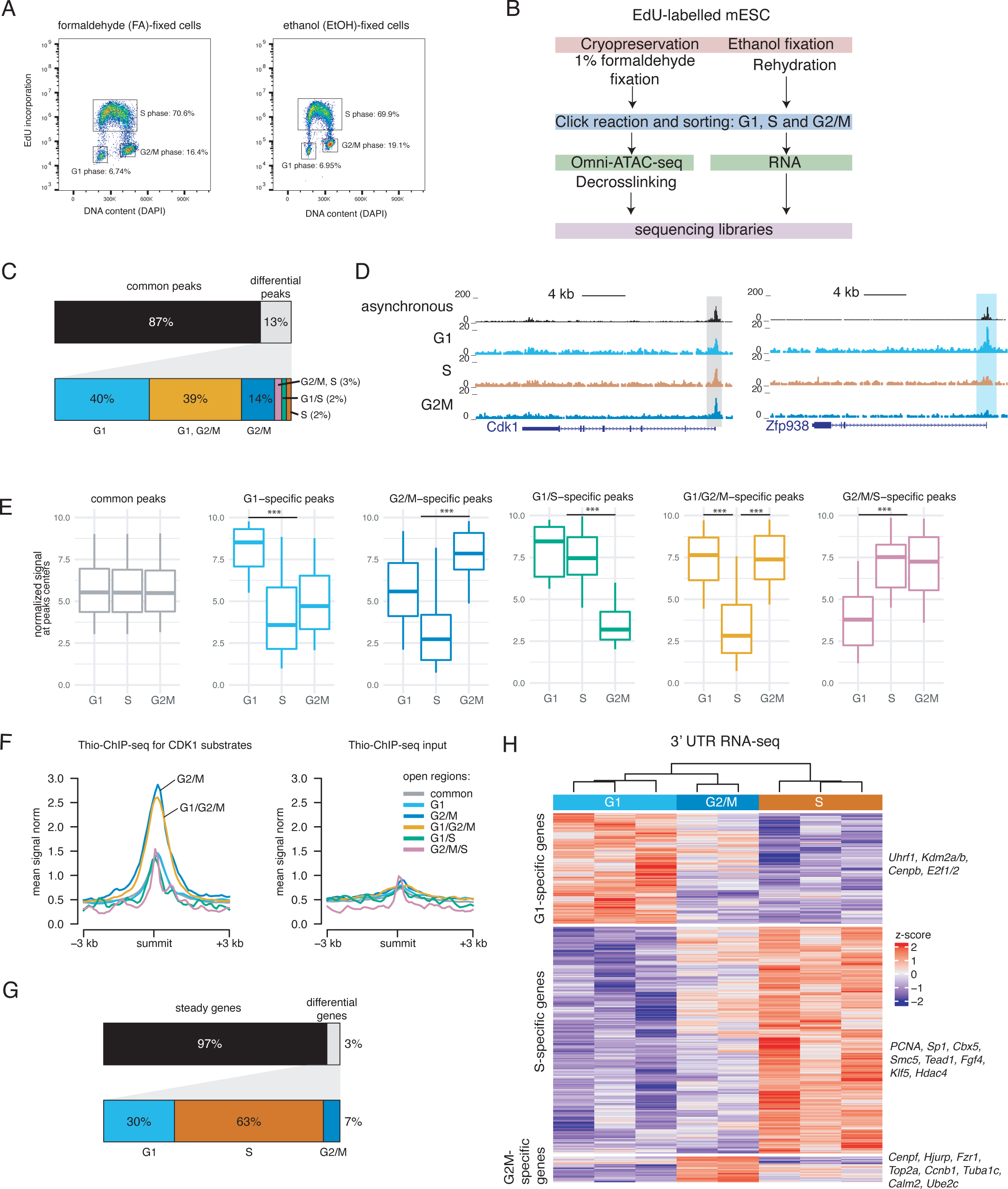
Cell cycle-specific profiles of chromatin accessibility and gene expression in unperturbed mESCs. (A) Representative flow cytometry profiles of EdU-labeled, DAPI stained mESCs, fixed with formaldehyde (left) or ethanol (left). (B) Schematic of the workflow. (C) Percentage of common and differentially accessible ATAC-seq peaks (top) with the latter category broken into cell cycle-specific groups (bottom). A differential peak is defined as a peak with an increase in openness by ≥1.5-fold (padj:.:0.05, estimated based on two biological replicates) in comparison to one or two other phases. (D) UCSC genome browser screenshots of asynchronous, G1, S and G2/M ATAC-seq accessibility tracks. A common peak at *Cdk1* promoter highlighted in grey (left) and G1-specific peak at *Zfp938* – in blue (right). (E) Box plots showing the mean ATAC-seq signal for covered bases in peak summits (± 100 bp) for different categories. (***p-value < 2.2e-16). The box designates the 25^th^ and 75^th^ percentiles and is divided by the median; the whiskers extend to the 5^th^ and 95^th^ percentiles. (F) Average profiles of Thio-ChIP-seq and its input (Michowski et al. 2020) over different classes of cell cycle-specific open regions (± 3 kb). (G) Percentage of steady and differentially expressed genes (top) with the latter category broken into cell cycle-specific classes (bottom). A differential gene is defined as a gene with an increase in expression (padj:.:0.05, estimated based on two or three biological replicates) in comparison to one or two other phases. (H) Heatmap showing three classes of cycling genes based on hierarchical clustering of differential transcripts (see Methods for details). Known cell cycle-regulated and other interesting transcripts are listed on the right.

To assess chromatin accessibility in G1, S, and G2/M phases, we performed Omni-ATAC-seq (Corces et al., 2017) on FA-fixed, stained and sorted cells (Figure 1A,B). We recovered a representative set of high confidence peaks based on read counts and reproducibility between replicates. 87% (n=10513) of these peaks did not show any variation in accessibility in all three cell cycle phases (Figure 1D), while 13% (n=1569) did (fold change≥1.5, padjý0.05) (Figure 1 C, D, E, Figure S1F). Most of the dynamic peaks had increased accessibility in G1 (40%, n=623) or G1/G2/M phases (39%, n=619), whereas highly accessible peaks only associated with the S phase were underrepresented (<7% [n=104] for all related categories) (Figure 1C, D, E).

To evaluate the specificity of identified cell cycle-dependent open regions, we used published data on DNA binding of CDK1 substrates in mESCs identified by thio-chromatin immunoprecipitation sequencing (Thio-ChIP-seq) (Michowski et al., 2020). By calculating average profiles for Thio-ChIP-seq or its input over common and differentially open regions, we observed low and comparable signals for all categories of peaks, but G2/M- and G1/G2/M-specific loci (Figure 1F). As expected, CDK1 substrates showed a 1.6-fold increase in binding over regions with elevated accessibility during G2/M progression or entry into G1. This suggests that identified dynamically accessible regions are regulated by cell cycle-dependent kinases, such as CDK1.

We then performed a 3’ mRNA-seq that allowed us to quantify levels of transcripts in fixed samples with sub-optimal quality and from a limited amount of material. In total, we detected only ∼3% of expressed genes (CPM ≥ 0.08 in asynchronous cells) to be differentially regulated in any of the cell cycle transitions (padj ý 0.05) (Figure 1G, H). The median fold change, however, was only ∼1.5-fold, indicating relatively mild overall changes in transcript levels. Gene set enrichment analysis (GSEA) for gene ontology (GO) terms showed that the differentially expressed genes were enriched for terms related to the cell cycle, its regulation and G2 to M transition (Figure S1G). Many of the differentially expressed genes (Figure 1H) are regulated by the E2F transcription factors and have an established function in specific cell cycle phases. More specifically, G2/M genes included a known topoisomerase, cyclin and members of the anaphase-promoting complex/cyclosome (APC/C) (*Top2a*, *Ube2c, Tuba1c, Calm2, Fzr1, Ccnb1, Tuba1c)*. Genes induced at the G1/S transition, DNA methylation maintenance and replication (*E2f1* and *2, Cenpb, Uhrf1* and *PCNA*) were among G1- and S-specific genes (Figure 1H). Interestingly, we also identified genes encoding important chromatin modifiers (*Hdac4, Kdm2a* and *b*) among the differentially expressed genes.

Taken together, we successfully combined ATAC-seq and RNA-seq with EdU-labeling and staining of fixed cells followed by sorting to obtain chromatin accessibility maps and transcriptional profiles in G1, S and G2/M phases in unperturbed mESC. We confirmed that genomic loci with increased openness in G2/M phase were enriched for binding of CDK1 substrates and we recovered known differentially expressed transcripts. Thus, we have established a flexible and robust method for assessing chromatin and transcriptional states on fixed cells during the cell cycle.

### Pharmacological synchronization of cells leads to aberrant chromatin accessibility and gene expression

To assess the impact of pharmacological inhibition on chromatin accessibility and gene expression, we performed omniATAC-seq and RNA-seq in mESC synchronized with the three most commonly used inhibitors (thymidine [G1], nocodazole [M] and CDK1i [G2/M]) (Figure 2A). We then compared the observed changes to the ones detected in sorted cell cycle populations.

**Figure 2.**
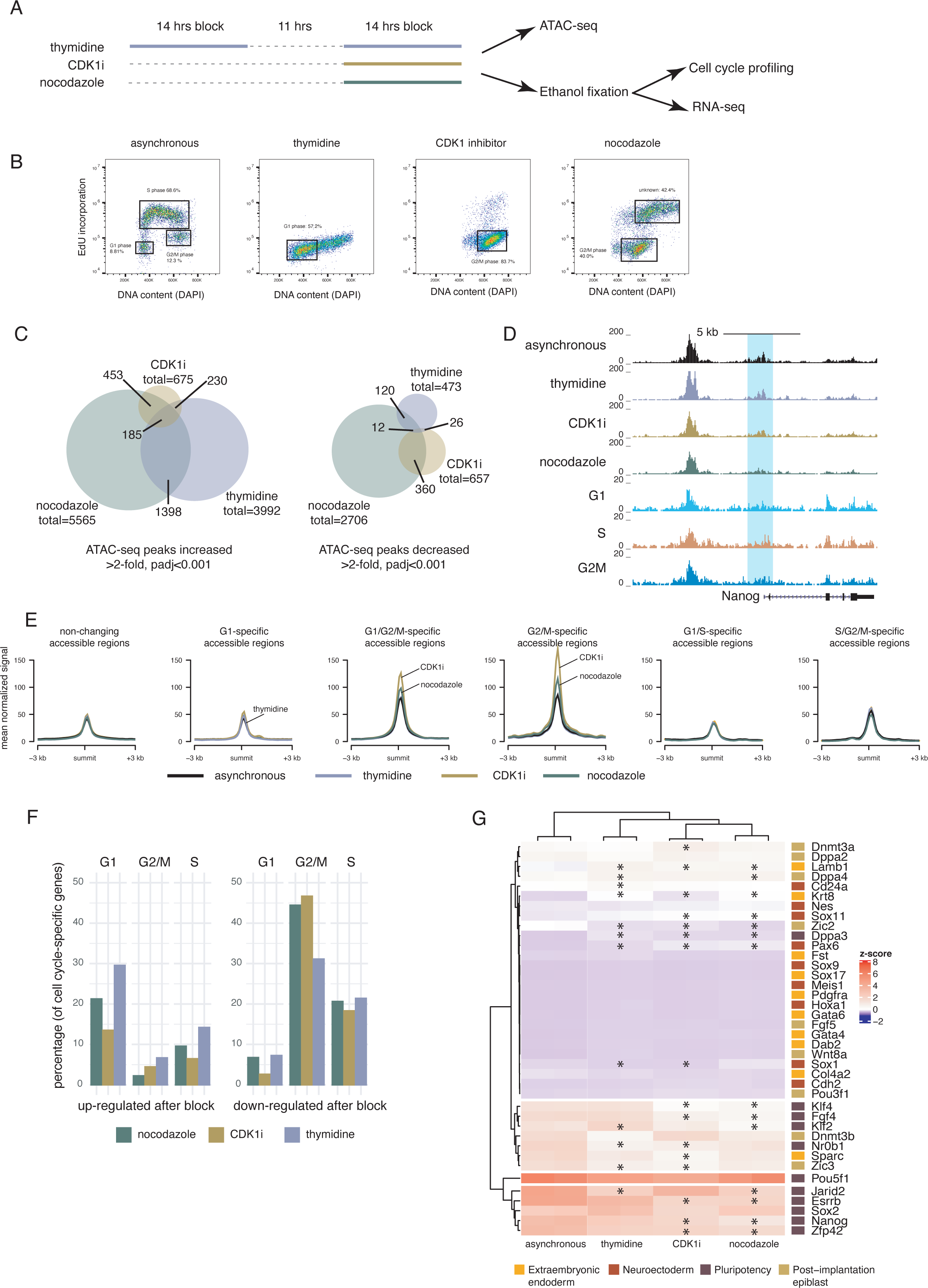
Impact of pharmacological synchronization on chromatin accessibility and gene expression. (A) Schematic of the workflow. (B) Representative flow cytometry profiles of mESC synchronized with double thymidine block, CDK1 inhibitor or nocodazole. (C) Venn diagrams showing the overlap of up-regulated (left) or down-regulated (right) peaks after drug treatments. (D) UCSC genome browser screenshot of ATAC-seq tracks from asynchronous, thymidine-, CDKi- or nocodazole-treated cells as well as from G1, S and G2/M sorted populations of cells. The peak at *Nanog* promoter is highlighted in blue. (E) Average profiles of ATAC-seq signal from asynchronous, thymidine-, CDKi- or nocodazole-treated cells over different classes of identified cell cycle-specific open regions (± 3 kb) in unperturbed mESC. (F) Bar plots showing the percentage of cycling genes identified in unperturbed mESC that were up- or down-regulated (fold change >1.5, padj<0.05) in mESC treated with thymidine, CDKi or nocodazole. (G) Heatmap showing mRNA steady levels of pluripotency and differentiation genes in asynchronous, thymidine-, CDKi- or nocodazole-treated cells (two biological replicates each). The genes were clustered based on their expression levels. The right panel indicates their class and names. *: fold change >1.5, padj<0.05.

We determined the efficiency of the blocks by EdU and DAPI staining as well as co-staining for H3S10phosho, a mitotic marker. Double thymidine block in mESC resulted in an accumulation of 57% cells in G1 phase (Figure 2B) and the rest remained in G2 ( Figure S2A) similar to the thymidine synchronization of hPSCs (Yiangou et al., 2019). 40% of cells blocked with nocodazole accumulated in G2/M, whereas the rest appeared in the late S and areas of higher DNA content (Figure 2B), suggesting endoreplication. Although we observed a significant enrichment of cells in M phase, there was still a large G2 fraction (Figure S2A). Thus, we will refer to the nocodazole synchronized population as G2/M phase. CDK1 inhibition successfully enriched 83% of cells in G2/M (Figure 2B) with most of the cells being at the G2-to-M border (Figure S2A).

The pharmacological treatments led to changes in chromatin accessibility in comparison to asynchronous mESCs, with nocodazole evoking the most prominent alterations (8271 differential peaks) and CDK1i - the least (1332 differential peaks) (Figure 2C, D, Figure S2B). Only 5.4% of increased and 0.9% of decreased peaks were shared for all conditions. The overlap for CDK1i- and nocodazole-treated cells was 14.5% and 21.4%; and higher than the one observed by comparing CDK1i vs thymidine treatment (4.8% for increased and 4.6% for decreased peaks) (Figure 2C). Interestingly, considerably more regions increased their accessibility than decreased upon nocodazole and thymidine synchronization (2-times and 8.4-times more, respectively); while up- and down-regulation were more balanced for CDK1i-treated mESCs (Figure 2C).

We next compared changes in accessibility induced by cell cycle inhibition in regions with the *bona fide* cell cycle-related changes as defined by ATAC-seq in the sorted populations. Thymidine block did not lead to any changes compared to asynchronous cells, whereas CDK1i and nocodazole induced significant opening of G2/M and G1/G2/M-specific regions, but not in others (Figure 2E). Together, these results imply that CDK1i has a mild effect on chromatin compaction, and similar to nocodazole it evoked changes in accessibility specific to G2/M open loci.

Unlike biased changes in ATAC-seq towards increased accessibility, comparable numbers of genes were up- and down-regulated upon synchronization (fold change >1.5, padj<0.05) (Figure S2B). Despite distinct responses to the inhibitors on the level of chromatin compaction and DNA content (Figure 2B, C), ∼43% of up- or down-regulated transcripts were shared for all conditions (Figure S2B). To elucidate what types of genes became deregulated, we performed gene set enrichment analysis for gene ontology terms. CDK1i and nocodazole treatment affected the expression of genes associated with cell cycle and mitosis, whereas the thymidine block up-regulated nuclear-encoded genes required for mitochondrial function and structure (Figure S2D). Both cytosolic (TK1) and mitochondrial (TK2) tyrosine kinases showed 2-fold up-regulation (padj<0.01) as well as mitochondrial pyrimidine nucleotides’ transporter *Slc25a36* and several oxidoreductases. Due to the limited number of reports that explored the link between the cell cycle and mitochondrial function, we can only speculate that inhibition with thymidine led to overcompensation of mitochondrial function due to an imbalance in the cytosolic dNTP pool (Pontarin et al., 2003).

To examine the effect of synchronization on cell cycle-dependent genes, we calculated the proportion of G1-, S- or G2/M-specific transcripts that were up- or down-regulated upon inhibition. The largest fraction (29.5%) of increased genes was G1-specific genes that responded to the thymidine treatment, suggesting partial activation of G1 phase program. In contrast, 31.1 - 46.7% of G2/M-specific genes (*Cenpf*, *Top2a*, *Ccnb1*, *Ube2c*, etc.) had a drop in their expression levels after all three types of blocks, indicating that synchronized cells did not acquire the transcriptional properties of G2/M phase. While thymidine-treated cells were expected to have a decrease in expression of G2/M-specific genes, the reduction observed in nocodazole and CDK1i synchronized cells could be caused by the prolonged G2/M arrest.

To investigate how synchronization affected the pluripotency state of mESC, we compared mRNA expression levels of genes associated with naïve state (*Nanog, Pou5f1, Sox2*, etc.) as well as differentiation markers for various lineages (*Fgf5, Sox1, Meis1*, etc.). Most of the lineage-specific markers were not present or lowly expressed in asynchronous mESC as well as in synchronized cells (Figure 2G). However, inhibition with CDK1i or nocodazole led to down-regulation (>1.5-fold, padj<0.05) of many pluripotency factors including *Nanog*, *Zfp42* (*Rex1*), *Klf4* and *Fgf4*, but not of the highly expressed *Pou5f1* (Figure 2G). Interestingly, the decrease in *Nanog* expression correlated with the decrease in chromatin accessibility at its promoter after synchronization (Figure 2D). This suggests that inhibition of cell cycle progression does not lead to spontaneous differentiation, but down- regulation of naïve markers, potentially making cells more susceptible to lineage-specifying cues.

We further investigated the inhibitors’ effects on mESC stemness by performing a colony-forming assay after releasing the cells from the blocks (Figure S3A, B). Unlike nocodazole-treated cells, thymidine- and CDK1i-blocked mESCs recovered and formed multiple colonies comparable to asynchronous control (Figure S3B). It indicates that the inhibitors partially or fully affect cell recovery, attachment or proliferation capacities.

To explore which factors could promote increased chromatin accessibility, we performed a motif analysis of differential ATAC-seq peaks [36]. Surprisingly, the motif associated with the p53 proteins family (p53, p73 and p63) was enriched in up-regulated peaks upon all types of blocks (Figure S3C, D, E, F). P53 and p73 transcripts were expressed in mESC and p53 protein was stabilized, especially upon nocodazole treatment (Figure S3G). It suggests that the increased accessibility can be at least partially due to the excessive activation of p53 family proteins.

Taken together, we have shown that pharmacological inhibition affected chromatin and gene expression status to a varying degree. Nocodazole and CDK1i treatment led to changes that resembled G2/M-specific chromatin state and DNA content, however, resulted in down-regulation of the G2/M-specific transcriptional program. In contrast, double thymidine block resulted in G1-like DNA content synchronization and partial activation of G1-specific genes, whereas it erroneously up-regulated nuclear-encoded mitochondrial genes and was insufficient to increase the accessibility of G1-specific open regions.

### Cell cycle-dependent chromatin accessibility changes correlate with specific chromatin signatures

To determine whether the cell cycle-specific ATAC peaks were associated with distinct chromatin signatures, we first compared the peaks to an annotation of mESCs chromatin states across the genome (Pintacuda et al., 2017). We used the chromHMM model, which provided a systematic annotation of genomic elements and was based on combination of marks characteristic to promoters (H3K4me3, H3K9ac, H3K27ac, NANOG, POU5f1), enhancers (H3K4me1, H3K27ac, H3K9ac, H3K27ac, NANOG, POU5f1), insulator sites (CTCF) and other states (Pintacuda et al., 2017). Regions stably open throughout the cell cycle were enriched for categories defined as active promoters (33.2% vs 5.6% of all chromHMM categories) and bivalent chromatin (6.9% vs 3.8% of all chromHMM categories) (Figure 3A, Figure S4A). Although common peaks overlapped enhancers (22.3%), insulators (8.3%), and intergenic regions (23.8%), their distributions did not largely differ from the genome-wide frequencies of chromHMM states (Figure 3A, Figure S4A). G2/M and G1/G2/M peaks mostly resembled common peaks in their chromatin signatures, whereas G1-specific open regions were enriched for insulators (1.5-fold; 13% vs 8.8% for all chromHMM categories) (Figure 3A, B, Figure S4A). We then confirmed CTCF motif enrichment in G1-specific peaks (Leporcq et al., 2020) (4.6- fold, p-value<10^-13^; Figure S4B) and its binding to a subset of G1 peaks (Stadler et al., 2011) (Figure 3B).

**Figure 3.**
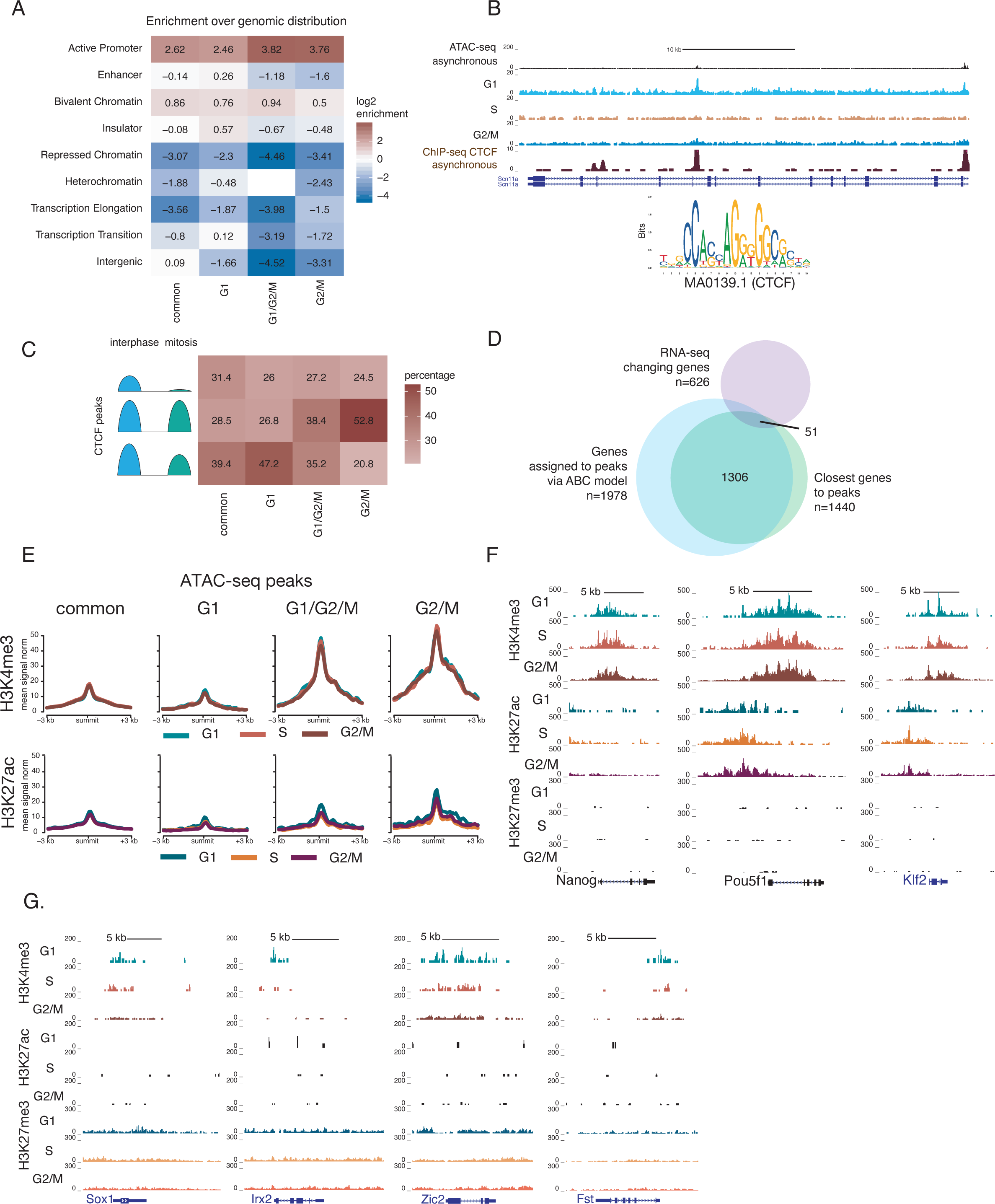
Chromatin states of cell cycle-dependent accessible regions and genes. (A) Heatmap showing log2-fold enrichment of co-occurrences for common or cell cycle-specific ATAC peaks and chromatin states (chromHMM) (Pintacuda et al., 2017). (B) UCSC genome browser screenshot of ATAC-seq tracks for asynchronous, G1, S and G2/M cells as well as CTCF ChIP-seq in asynchronous mESCs (Stadler et al., 2011). The sequence logo shows the CTCF motif enriched in ATAC peaks. (C) Heatmap showing the percentage of CTCF-bound cell cycle-dependent ATAC-seq belonging to different categories of CTCF binding defined in (Owens et al., 2019). (D) Venn diagram showing the overlap between *bona fide* cell-cycle genes, closest genes to cell cycle-specific open regions and genes assigned to these regions via ABC model. (E) Average profiles over different categories of ATAC-seq peaks (± 3 kb) for H3K4me3 and H3K27ac in G1, S and G2/M cells sorted from unperturbed mESC. H3K27me3 profiles are in Figure S4F. (F) (G) UCSC genome browser screenshots of H3K4me3, H3K27ac and H3K27me3 signals for G1, S and G2/M cells sorted from unperturbed mESC. Panel (F) shows pluripotency genes and panel (G) - lineage-specific genes that are transiently bivalent during the cell cycle.

To further explore the link between the cell cycle-specific changes in chromatin accessibility and CTCF binding, we took advantage of published CTCF ChIP-seq data generated for interphase and mitotic mESCs (Owens et al., 2019). CTCF mitotic bookmarking has been reported to exist in mESCs and not in somatic cell lines (Owens et al., 2019). 47.2% of CTCF-bound G1-specific open regions were strongly bound by CTCF in interphase with reduced binding in mitosis (Figure 3C). In contrast, only 26% and 26.8% of peaks overlapped other categories of bookmarking regions (lost in mitosis and similar in both phases as defined in Owens et al.) (Figure 3C). Interestingly, a large fraction of CTCF-bound G2/M accessible regions (52.8%) fell into a category of bookmarked regions that did not significantly change CTCF enrichment between interphase and mitosis (Figure 3C). The results indicate that cell cycle-dependent loci have specific chromatin signatures (of promoters, bivalent chromatin and insulators) and potentially different functional roles, including structural organization of chromatin by CTCF in G2/M to G1 transition.

Unlike GO terms for cell cycle-regulated genes identified by RNA-seq (Figure 1H, Figure S1G), a similar analysis for genes with promoters overlapping or near differential ATAC-seq peaks (padj≤0.05) did not show enrichment for categories identified by RNA-seq (Figure 1H, Figure S1G), a similar analysis for genes with promoters overlapping or near differential ATAC-seq peaks (padj≤0.05) did not show enrichment for categories related to the cell cycle ( Figure S3C). 60.1% of changing ATAC-seq peaks (padj≤0.05) were promoter-distal, making the peak-to-gene assignment ambiguous. To examine if erroneous misassignments could mask the effect on the cell cycle genes, we performed the analysis for only differentially accessible promoters and also assigned distal open regions to promoters through the activity-by-contact (ABC) model (Fulco et al., 2019). We built the ABC model (Fulco et al., 2019) to predict the genome-wide map of enhancer-gene pairs by incorporating H3K27ac (Atlasi et al., 2019), *in situ* Hi-C (Yan et al., 2018) and gene expression data from mESCs grown in serum/LIF. Although the ABC model allowed us to expand the number of putative targets from 1440 to 1978, the overlap with cell cycle-regulated genes remained modest (Figure 3D) and did not substantially change GSEA analysis (Figure S4C). These results show that differential expression of genes during the cell cycle does not correlate with changes in chromatin accessibility, indicating that they cannot be explained by these alterations.

To assess whether the changes in gene expression and accessibility correlated with changes in histone modifications, we performed CUT&Tag (Kaya-Okur et al., 2019) on sorted cell cycle phases for H3K4me3, H3K27ac and H3K27me3 marks. This minimal set of chromatin marks allowed us to define putative promoters, enhancers and repressed domains. As expected, the promoters of the cycling genes (as defined by <u>C</u>ap <u>A</u>nalysis of <u>G</u>ene <u>E</u>xpression [CAGE] (Lizio et al., 2019)) were depleted for H3K27me3 ( Figure S4F), however, they had high levels of H3K4me3 throughout the cell cycle (Figure S4D, E). The cycling genes were also associated with low amounts of H3K27ac throughout the cell cycle. However, we found an even lower enrichment of H3K27ac was found in G1 phase (Figure S4D, E). ATAC-seq peaks with increased accessibility in G2/M- and G1/G2/M-specific were especially enriched for active promoters (Figure 3A, Figure S4A), and accordingly had high but stable H3K4me3 signal in distinct cell cycle phases (Figure 3E). H3K27 acetylation and trimethylation did not show any significant changes either (Figure 3E, Figure S4F).

Several reports have linked propensity towards pluripotency or differentiation with cell cycle phases in hESCs (Gonzales et al., 2015; L. Liu et al., 2019; Y. Liu et al., 2017; Singh et al., 2013; 2015) and mouse cells. Promoters and distal regulatory elements of pluripotency regulators had stable H3K4 trimethylation as well as H3K27 acetylation (Figure 3F), and their transcript levels did not change during the cell cycle. Unbiased genome-wide analysis, as well as analysis focused on selected groups of regulatory regions, did not identify genomic loci with significant changes in chromatin modifications. Nevertheless, we found individual domains with moderate G1-specific levels of H3K4me3 that were absent in other phases (Figure 3G). Some of these domains were also marked by stable levels of H3K27me3 and located near neuroectodermal (*Sox1*, *Irx2* and *Zic2*) and endodermal (*Fst*) genes (Figure 3G) with no or stable expression in all cell cycle phases. The fact that H3K4me3 is associated with these genes in G1 confirms a possible mechanism previously suggested for hESCs (Singh et al., 2015) of how and why embryonic cells in G1 phase appear more prone to differentiation.

Taken together, our results show that in addition to being compatible with profiling of gene expression and chromatin accessibility, cell cycle-specific sorting can be combined with the detection of histone modifications. Interestingly, we found a potential link between G1-specific regions and CTCF binding. However, the differential expression of genes during the cell cycle cannot be explained by changes in histone modifications or chromatin accessibility.

### The combination of cell cycle-sorting and SLAM-seq shows the correlation between steady and nascent transcript levels and differences in mRNA stability

We reasoned that steady-state mRNA levels might not properly reflect the dynamics of gene expression, which could potentially exacerbate the correlations with chromatin accessibility changes. To more precisely measure the rate of mRNA synthesis, we performed metabolic RNA sequencing, SLAM-seq (Herzog et al., 2017). To our knowledge, 4-thiourindin labeling (s^4^U) combined with 3’UTR sequencing has not been previously integrated with EdU labeling or with fluorescence-activated cell sorting (FACS) of fixed cells. To confirm the compatibility of both labeling approaches we assessed T-to-C conversion rates in EdU samples with and without s^4^U incorporation and did not detect any spurious conversions (Figure 4A, B).

**Figure 4.**
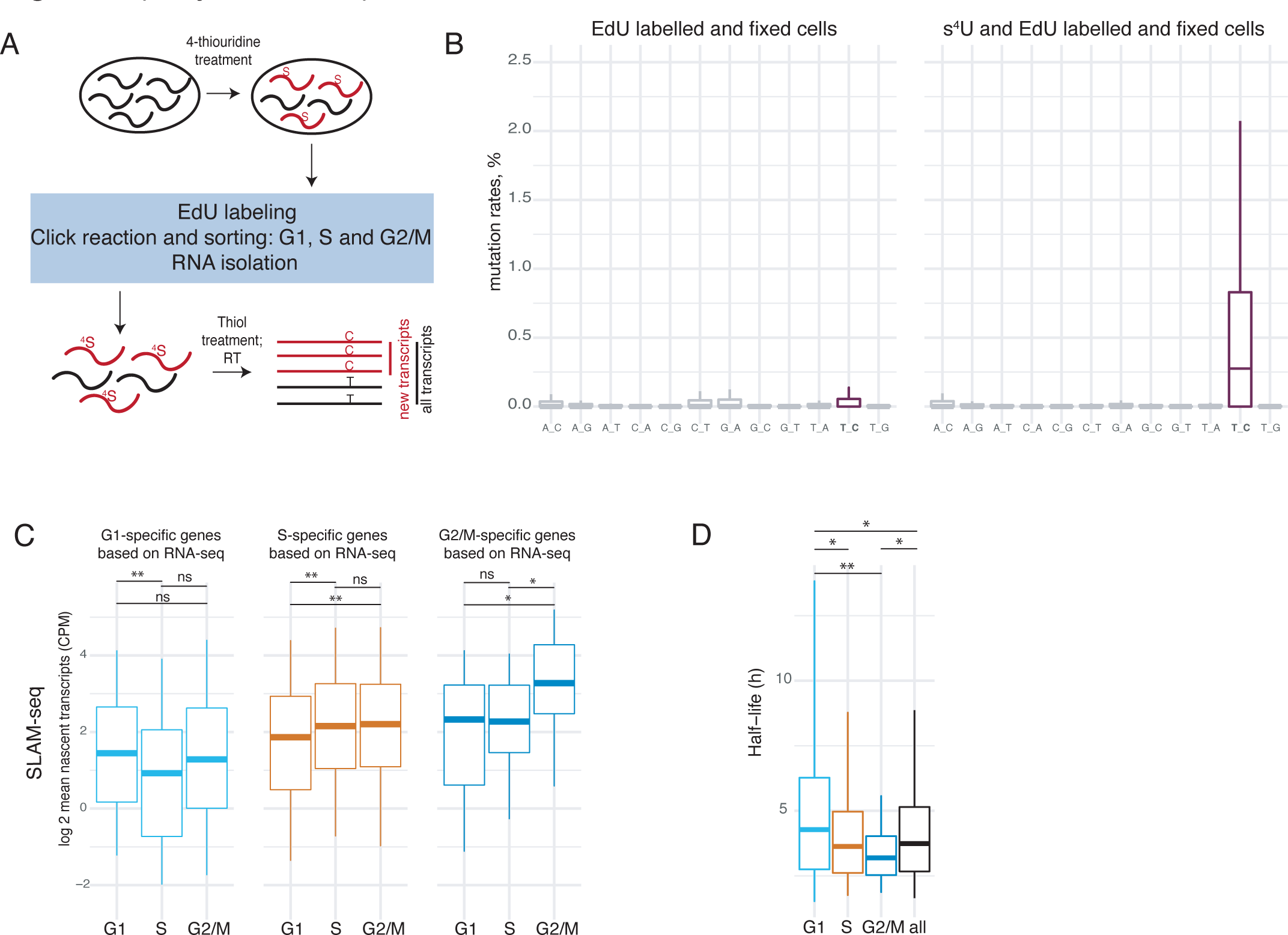
Detection of nascent transcription in individual cell cycle phases. (A) Schematic of the workflow for cell cycle-specific SLAM-seq. RT – reverse transcription. (B) Conversion rates in mapped reads, prepared from EdU-labelled mESCs without or with metabolic labeling (+ s^4^U). The box designates the 25^th^ and 75^th^ percentiles and is divided by the median; the whiskers extend to the 5^th^ and 95^th^ percentiles. (C) Box plots showing counts per million reads mapped (CPM, log2 scale) for nascent transcripts detected for G1-, S- or G2/M-specific genes. Wilcoxon rank-sum test: *p-value<0.001, **p- value<10^-5^, ns – not significant. (D) Box plots depict half-lives (h) of mESC transcripts (Herzog et al., 2017) broken into G1-, S- and G2/M-specific categories. Wilcoxon rank-sum test: *p-value<0.01, **p-value<0.05.

To assess whether changes in ATAC-seq led to any shift in steady or nascent transcripts levels, we compared counts per million reads mapped (CPM) for genes near cell cycle-dependent open regions. In agreement with our previous observations, there were no changes in median levels of steady state transcripts (Figure S5A) and alterations in the levels of nascent transcripts largely resembled the steady state (Figure S5B).

We then examined the relationship between cell cycle-specific genes based on steady state levels and their nascent transcript levels. Genes with higher expression in G1 phase based on 3’UTR RNA-seq had similar levels of nascent transcription in both G2/M and G1 phases, whereas S-phase specific genes had an increase in nascent transcription coinciding with S and G2/M. We observed the most prominent change (1.5-fold, pval<0.001) for G2/M-specific genes: the burst of nascent transcription occurred at late S and G2/M (Figure 4C). A recent study showed that there were different avenues for mRNA turnover during the cell cycle in human cells (Battich et al., 2020).

To address the mRNA turnover of the identified cycling genes, we compared their mRNA half-lives with pulse-chase SLAM-seq data in mESCs (Herzog et al., 2017). While S-specific genes largely resembled all expressed genes, G1-specific genes had longer half-lives than all expressed genes (4.3 h vs 3.7 h, p-value<0.05) and other categories of cycling transcripts. In contrast, the half-lives of G2/M-related transcripts were the shortest (3.2 h vs 3.7 h, p-value<0.05), indicating that a degradation mechanism might substantially contribute to the regulation of G2 mRNA.

Taken together, our data suggest that cell cycle-dependent mRNA levels could be regulated at the level of initiation of transcription and mRNA stability.

## DISCUSSION

In this study, we integrated cell cycle-specific sorting with major functional genomics techniques and compared the *bona fide* cell cycle-dependent transcriptional and chromatin states with the ones obtained from pharmacologically synchronized cells. Our results showed that synchronization with inhibitors caused changes in chromatin compaction and gene expression, which partially correspond to actual cell cycle-dependent alterations. Besides better synchronization of mESCs compared to thymidine and nocodazole, CDK1 inhibitor evoked milder overall changes in chromatin accessibility. Bioinformatics analysis suggests that CDK1 substrates and CTCF are possible mediators of cell cycle-specific variations in chromatin openness.

A combination of brief EdU-labeling with staining of DNA content is the least invasive and versatile approach to sort out specific cell cycle phases. Covalent binding of the fluorophore to the nascent DNA strand via click chemistry does not require harsh permeabilization and DNA denaturation, thus, it is compatible with chromatin assays (Stewart-Morgan et al., 2019), (Gutiérrez et al., 2019), (Kliszczak et al., 2011). It does not limit experiments to one reporter cell line and allows sufficient separation of S phase from G1 and G2/M. Being non-immune reactive, click chemistry can be combined with antibody-based co-stainings to detect membrane receptors, cellular and nuclear factors (e.g., H3S10phospho to label M phase), thereby expanding its use. We demonstrate the compatibility of the approach with aldehyde- and alcohol-based fixatives, thus, making it suitable for any downstream assays such as ATAC-seq, CUT&Tag and RNA-seq. Moreover, we have shown the feasibility of simultaneous labeling of DNA and nascent mRNA (Herzog et al., 2017), which allows estimation of the fraction of new transcripts in individual cell cycle phases.

At each point of the cell cycle, a cell has a distinct constellation of properties such as DNA content, chromatin and 3D landscapes, rate of transcription, proteomic and metabolic states. Pharmacological blocks result in cells aligned for such a characteristic as DNA content. Synchronization is meant to produce large enough quantities of cells to study events happening during specific phases of a normal unperturbed cell cycle. Although there are quite some cell cycle inhibitors (Yiangou et al., 2019), only a few are routinely used in the field of DNA replication (Francis et al., 2009; Reverón-Gómez et al., 2018) and mitosis (Y. Liu et al., 2017; H. Zhang et al., 2019). Moreover, there are no inhibitors to enrich for G2 phase alone, which makes G2-specific chromatin and transcriptional changes challenging to explore. In this study, we showed that treatments with thymidine and nocodazole led to considerable changes in chromatin accessibility but only moderately recapitulated the normal cell cycle phases. Interestingly, the CDK1 inhibitor evoked relatively mild changes in chromatin compaction that resembled G2/M phase. In contrast to variations in chromatin openness, only a subset of genes was appropriately up-regulated by the drug treatments, while a considerable fraction of G2/M-specific transcripts even showed down-regulation. This anti-synchronization of gene expression can be a prerequisite or consequence of prolonged resistance to the cell cycle progression. The large discrepancy between unperturbed cell cycle phases and arrested cells has also been reported for proteome in a human leukemia cell line (Ly et al., 2017; 2015).

Precisely regulated gene expression is key for controlling the progression through the cell cycle and successful cell division. By profiling chromatin and transcriptional states in unperturbed cells, we identified changes in chromatin accessibility, repressive and active marks as well as gene expression. Several genes with cell cycle-regulated expression have essential phase-specific roles during the cell cycle (Battich et al., 2020). Surprisingly, we did not identify changes in accessibility and histone posttranslational modifications at their promoters or their putative regulatory regions that we predicted from a comprehensive combination of functional genomics data including three-dimensional chromosomal structures (Fulco et al., 2019). Therefore, we speculate that regulation can occur at the level of differential recruitment of transcription factors and co-factors to constitutively open regions. However, we do not exclude that other chromatin-dependent mechanisms can play a role in transient activation of cell cycle-dependent transcripts or help G2-specific genes escape transient repression associated with dosage compensation in G2 phase (Skinner et al., 2016; Voichek et al., 2016).

The genomic regions, where we detected increased accessibility in G2/M phase, correlated with the binding of CDK1 substrates, which included many chromatin modifiers such as histone demethylases (Michowski et al., 2020). The diversity of putative targets which activity might depend on CDK1 (Kim et al., 2018; Michowski et al., 2020) can lead to very different functional outcomes at the level of gene regulation. We also showed that the G1-specific open regions were enriched for CTCF motif and their large fraction overlapped CTCF sites with higher binding in interphase (Owens et al., 2019). CTCF was shown to maintain local nucleosomal positioning for timely reactivation of gene expression after the passage of the replication fork and mitosis in mESCs (Owens et al., 2019). Our analysis implies that the transcriptional outcomes of these periodic changes in unstimulated cells are modest and could be properly buffered at the level of posttranscriptional regulation affecting mRNA and protein stability. Although retention of regulatory proteins on mitotic chromatin and its importance for re-initiation of transcription have been actively studied (Festuccia et al., 2019; Owens et al., 2019; Teves et al., 2016), the differences and functional consequences of binding in different cell cycle phases have also been gaining attention (Friman et al., 2019).

Our analysis suggests that transient emergence of G2-specific transcripts might rely on degradation to a greater extent than for other mRNAs, as G2-specific transcripts have the shortest half-lives when compared to all cycling genes. A study in yeast proposed that such degradation following a peak in mRNA expression of cycling genes was a passive process (Eser et al., 2014). Another study found and characterized a mechanism of mitosis-induced decay for some transcripts (Trcek et al., 2011). The most recent single-cell work on the cell cycle implied different strategies for synthesis and degradation dynamics for cycling genes and observed that functionally similar transcripts tended to utilize comparable strategies (Battich et al., 2020). However, the molecular mechanism has yet to be explored.

In summary, we have shown that synchronization of cells by EdU labeling followed by FACS sorting is the least invasive method to enrich cells in the different phases of the cell cycle. We demonstrate that this method is compatible with various genomics and transcriptomics techniques, allowing the determination of chromatin and transcriptional states with specific phases of the cell cycle. We also show that pharmacological inhibition leads to aberrant changes in chromatin accessibility but not transcription. We foresee that more genomics techniques will become compatible with a small number of fixed cells and more cell cycle-related biological questions will be addressed in unperturbed cells in various conditions and genetic backgrounds.

## METHODS

### Cell culture

E14 mouse ES cells (male, RRID: CVCL_C320) were cultured in the serum/LIF medium containing Glasgow Minimum Essential Media (Sigma), 10% heat-inactivated FBS (HyClone), 1x Penicillin-Streptomycin (Thermo Fisher Scientific), 2 mM GlutaMAX (Thermo Fisher Scientific), 100 µM β-Mercaptoethanol (Thermo Fisher Scientific), 1 mM sodium pyruvate (Thermo Fisher Scientific), 0.1 mM MEM Non-Essential Amino Acids Solution (Thermo Fisher Scientific) and Leukemia Inhibitory Factor (homemade). Cells were passaged by trypsinization (Sigma) every two days. The cells were routinely tested and were mycoplasma negative.

### Treatment with inhibitors

mESCs were seeded at 30-50% confluency 8 hours before the start of treatment. Cells were treated with 2 mM thymidine (Sigma, T1895) for 14 hr, washed with PBS and grown for 11 hrs before additional thymidine treatment for 14 hrs; 100 ng/ml nocodazole (Sigma, M1404) for 14 hr; 10 µM CDK1 inhibitor (Ro-3306, Selleckchem, S7747) for 14 hrs.

### Cell cycle phase sorting

After three washes with PBS, treated or non-treated cells were incubated in media supplemented with 20 nM EdU (Lumiprobe, 10540) for 10 min in the incubator, washed with PBS three times and trypsinized (Sigma). Cell pellets were additionally washed three times with PBS before fixation. Ethanol fixation (for flow cytometer analysis or sorting followed by RNA-seq or SLAM-seq) was done by drop-wise addition of 100% cold ethanol to cells resuspended in PBS so that the resulting concentration was 70% of ethanol. Cells were fixed and stored at -20°C for minimum 12 hours. mESC for omni-ATAC-seq and CUT&Tag were fixed in 1% formaldehyde (Thermo Fisher SCIENTIFIC) and quenched by adding glycine (final concentration 0.125M) (Sigma). Up to 5 million cells per reaction were extensively washed in PBS with 1% FBS. The cells were resuspended in 200 μl of 1x BD Perm/wash buffer (BD Biosciences, 554723) and incubated at room temperature for 15 min. Before Click reaction, the cells were washed in PBS with 1% FBS. Cell pellets were resuspended in 400 μl Click reaction [1x PBS, 2 mM CuSO4 (Sigma, 12849), 20 mg/ml L-ascorbic acid sodium salt (Acros Organics, AC352685000), 10 μM Sulfo-Cyanine5 azide (Lumiprobe, A3330)] and incubated in the dark at room temperature for 30 min. After two washes with 1% FBS in PBS, the cells were resuspended in 1x BD Perm/wash buffer (BD Biosciences, 554723) with 2.5 ng/μl DAPI (Sigma) and incubated for 1 hr at +4°C before the flow cytometry analysis (Beckman Coulter CytoFlex) or sorting (SONY MA900). For mitotic cells staining we incubated cells with 1:500 Anti-phospho-Histone H3 (Ser10) Antibody (Sigma, 06-570) for 1 hr at room temperature, followed by incubation (1 hour, room temperature) with 1:1000 Goat anti-Rabbit IgG (H+L) Highly Cross-Adsorbed Secondary Antibody, Alexa Fluor 488 (Thermofisher Scientific, A-11034).

### Omni-ATAC-seq on sorted cell cycle phases and synchronized cells

50,000 cells for each cell cycle phase (G1, S or G2/M) were sorted. The main component for all buffers was the resuspension buffer: 10 mM Tris-HCl, 10 mM NaCl, 3 mM MgCl2 in water. Cell pellets were resuspended in 50 μl of cold lysis buffer [resuspension buffer, 0.1% v/v NP-40 (Research Products International), 0.1% Tween-20 (Sigma), 0.01% v/v digitonin (Promega)] and incubated on ice for 3 min. After one wash with 1 ml wash buffer [resuspension buffer, 0.1% Tween-20] the cells were pelleted at 500 g for 10 min at +4°C. The cell pellet was transposed in 50 μl of the following reaction mix: 1x tagmentation buffer (Illumina, 20034197), 1x PBS, 0.1% v/v Tween-20, 0.01% v/v digitonin, 2.5 μl of tagment DNA enzyme 1 (Illumina, 20034197), water. Cells were incubated in the thermomixer (1000 rpm) at +37°C for 30 min. Reverse crosslinking was done by adding 200 μl of the decrosslinking solution [50 mM Tris-Cl, pH 7.5, 1 mM EDTA, 1% SDS, 0.2M NaCl, 5 ng/ml Proteinase K (Thermo Fisher Scientific)] and incubating the cells at +65°C at 1000 rpm in the thermomixer overnight. Tagmented DNA was purified with MinElute PCR Purification Kit (QIAGEN, 28004) and eluted in 10 μl QIAGEN EB buffer. The sequencing libraries were amplified and barcoded with NEBNext® High-Fidelity 2X PCR Master Mix (NEB, M0541S) and size selected with AMPure XP beads (Beckman, A63880). All libraries were sequenced on NextSeq 500.

### CUT&Tag on sorted cell cycle phases

70,000 cells for each cell cycle phase (G1, S or G2/M) were sorted. We followed the published protocol (Kaya-Okur et al., 2019). The following antibodies were used: 1:100 anti-H3K4me3 (Cell Signaling, 9751), 1:100 anti-H3K27me3 (Cell signaling, 9733), 1:100 anti-H3K27ac [EP16602] (Abcam, ab177178), Guinea Pig anti-rabbit IgG (Antibodies-Online, ABIN101961). The permeabilized cells were incubated with the primary antibody overnight at +4°C and 30 min at room temperature with the secondary. The reverse crosslinking was performed at +65°C for 4 hrs by supplementing the tagmentation buffer with 20 mM EDTA, 0.2% SDS, 20 ng/ml Proteinase K (Thermo Fisher Scientific). Tagmented DNA was purified with MinElute PCR Purification Kit (QIAGEN, 28004) and eluted in 32 μl QIAGEN EB buffer. The sequencing libraries were amplified and barcoded with NEBNext® High-Fidelity 2X PCR Master Mix (NEB, M0541S) and size selected with AMPure XP beads (Beckman, A63880). All libraries were sequenced on NextSeq 500.

### RNA-seq, 3’UTR mRNA-seq combined with SLAM-seq

For SLAM-seq mESCs were incubated with 100 μM 4-thiouridine (Sigma, T4509) for 2 hrs before EdU labeling. After cell cycle phase sorting, RNA was extracted with QIAzol (QIAGEN) following the manufacturer’s instructions and the detailed SLAM-seq protocol (Herzog et al., 2020). 0.1 mM DTT was added during isopropanol precipitation and resuspension in water. Labeled RNA was protected from light. 500 ng of RNA underwent the iodoacetamide treatment [10 mM iodoacetamide, 50 mM NaPO4, pH 8.0, 50% DMSO] in 50 μl reaction. RNA-seq libraries were prepared with QuantSeq 3′ mRNA-Seq Library Prep Kit FWD for Illumina (Lexogen, 015.24). RNA from fixed synchronized cells or unperturbed mESCs was extracted with RNeasy Mini Kit (QIAGEN, 74104) and processed with Illumina TruSeq v2 kit (Illumina, RS-122-2001). All libraries were sequenced on NextSeq 500.

### Alkaline staining and Western blot

After treatment with inhibitors and trypsinization, mESCs were washed twice in PBS and counted. 1x10^3^ cells were seeded into a 6-well plate and cultured for 5 days. Cells were fixed in citrate-acetone-formaldehyde and stained using Alkaline Phosphatase Detection Kit (Sigma, SCR004). The rest of the cells were lysed in RIPA buffer (150 mM NaCl, 1% NP-40, 0.5% sodium deoxycholate, 0.1% SDS, 50 mM Tris, pH 8.0) and sonicated for 5 min (30 sec ON/ 30 sec off) with Bioruptor (Diagenode). Total protein was measured with a Bradford assay (Bio-Rad, 5000001), denatured in 2x Laemmli buffer (5 min at +95°C) and 20 µg were loaded per well of Mini-PROTEAN Precast Gel (Bio-Rad). The protein transfer was done with the Trans-Blot Turbo system (Bio-Rad). The membranes were incubated with 1:5 home-made anti-p53 (overnight at +4°C) and 1:100,000 anti-Lamin B1 (1 hr at room temperature) (Cell Signaling Technology, #12586). As secondary antibodies, we used IRDye 800CW Goat anti-Rabbit IgG and IRDye 680RD Goat anti-Mouse IgG (LI-COR) (1:1000). The membranes were imaged using Image Studio Lite software (Li-COR Biosciences).

### Data analysis for Omni-ATAC-seq and CUT&Tag

The reads were trimmed with cutadapt (Martin, 2011) in the pair-end mode and default parameters. After alignment with bowtie2 (Langmead and Salzberg, 2012) to mm10; reads were filtered (-q 10) and duplicates were removed with Picard tools. We called ATAC-seq peaks with MACS2 (Y. Zhang et al., 2008) on individual samples and combined all peaks to count reads over the union of all peaks with bamCoverage (deepTools (Ramírez et al., 2016)) or over CAGE-defined p1 and p2 TSSs (Lizio et al., 2019). The read counts were normalized and further processed with DESeq2 (Love et al., 2014). The regions were classified based on DESeq2 fold-changes and padj. The tracks were visualized in the UCSC Genome browser (Kent et al., 2002).

### Data analysis for 3’UTR mRNA-seq and SLAM-seq

SLAM-seq data combined with 3’UTR mRNA-seq were aligned with SLAMDUNK (slamdunk all -5 12 -n 100 -t 3 -m -rl 75) (Neumann et al., 2019). Mutational rates were obtained with alleyoop utrrates (-l 75). ReadCounts and TcCounts outputs were used for the analysis and normalized with DESeq2 (Love et al., 2014). The clusters of co-regulated genes were obtained with hierarchical cluster analysis (complete linkage clustering) of differentially expressed genes. The number of clusters (k=3) was selected based on the dendrogram and visualized with a complex heatmap package (Gu et al., 2016).

### Data analysis for RNA-seq

The reads were trimmed with cutadapt 1.15 (Martin, 2011) in the single-end mode (--quality 15) and aligned with STAR (default parameters) to mm10 followed by filtering with samtools (-q 10). Aligned reads were counted with HT-seq over Genecode vM17 annotation, normalized and further processed with DESeq2 (Love et al., 2014).

### Motif analysis and GSEA

The regions of interest (in bed format) were analyzed with TFmotifView (Leporcq et al., 2020) where they were matched to the non-redundant JASPAR CORE 2020 database of motifs. Fold change and significance were calculated based on genomic controls with matching GC content. GSEA was done on ranked transcripts based on p-value with msigdbr 7.2.1 and fgsea 1.14.0 R packages.

### Activity-by-contact model

We predicted enhancer-gene pairs by using the activity-by-contact model (Fulco et al., 2019) with the following input data for mESCs: ATAC-seq (this study), RNA-seq (this study), H3K27ac (SRR5110896 and SRR5110902) (Atlasi et al., 2019) and *in situ* Hi-C (4DNESU4BQU4G) (Yan et al., 2018). We used the default threshold (0.02) to obtain significant ABC scores and contacts (excluding promoter elements).

## DATA ACCESS

All raw sequencing data generated in this study can be accessed in the NCBI Sequence Read Archive (SRA; https://www.ncbi.nlm.nih.gov/sra) under accession number GSE167963.

## ACKNOWLEDGMENTS

We thank Kathleen Stewart-Morgan for the EdU labeling protocol, Tugce Aktas for the comments on the manuscript and the Fragile Nucleosome community. D.S. was in part funded by the European Union’s Horizon 2020 research and innovation program under the Marie Sklodowska- Curie grant agreement 749362. The work in the Helin through the Memorial Sloan Kettering Cancer Center Support Grant (NIH P30 CA008748) and through the CRUK Grand Challenge Programme (C5836/A29059)

## REFERENCES

Abramo, K., Valton, A.-L., Venev, S.V., Ozadam, H., Fox, A.N., Dekker, J., 2019. A chromosome folding intermediate at the condensin-to-cohesin transition during telophase. Nat. Cell Biol. 21, 1393–1402. doi:10.1038/s41556-019-0406-2

Atlasi, Y., Megchelenbrink, W., Peng, T., Habibi, E., Joshi, O., Wang, S.-Y., Wang, C., Logie, C., Poser, I., Marks, H., Stunnenberg, H.G., 2019. Epigenetic modulation of a hardwired 3D chromatin landscape in two naive states of pluripotency. Nat. Cell Biol. 21, 568–578. doi:10.1038/s41556-019-0310-9

Battich, N., Beumer, J., de Barbanson, B., Krenning, L., Baron, C.S., Tanenbaum, M.E., Clevers, H., van Oudenaarden, A., 2020. Sequencing metabolically labeled transcripts in single cells reveals mRNA turnover strategies. Science 367, 1151–1156. doi:10.1126/science.aax3072

Behera, V., Stonestrom, A.J., Hamagami, N., Hsiung, C.C., Keller, C.A., Giardine, B., Sidoli, S., Yuan, Z.-F., Bhanu, N.V., Werner, M.T., Wang, H., Garcia, B.A., Hardison, R.C., Blobel, G.A., 2019. Interrogating Histone Acetylation and BRD4 as Mitotic Bookmarks of Transcription. Cell Rep 27, 400–415.e5. doi:10.1016/j.celrep.2019.03.057

Bjursell, G., Reichard, P., 1973. Effects of thymidine on deoxyribonucleoside triphosphate pools and deoxyribonucleic acid synthesis in Chinese hamster ovary cells. J Biol Chem 248, 3904– 3909.

Cooper, S., Iyer, G., Tarquini, M., Bissett, P., 2006. Nocodazole does not synchronize cells: implications for cell-cycle control and whole-culture synchronization. Cell Tissue Res. 324, 237–242. doi:10.1007/s00441-005-0118-8

Corces, M.R., Trevino, A.E., Hamilton, E.G., Greenside, P.G., Sinnott-Armstrong, N.A., Vesuna, S., Satpathy, A.T., Rubin, A.J., Montine, K.S., Wu, B., Kathiria, A., Cho, S.W., Mumbach, M.R., Carter, A.C., Kasowski, M., Orloff, L.A., Risca, V.I., Kundaje, A., Khavari, P.A., Montine, T.J., Greenleaf, W.J., Chang, H.Y., 2017. An improved ATAC-seq protocol reduces background and enables interrogation of frozen tissues. Nat Methods 14, 959–962. doi:10.1038/nmeth.4396

Darzynkiewicz, Z., Traganos, F., Zhao, H., Halicka, H.D., Li, J., 2011. Cytometry of DNA replication and RNA synthesis: Historical perspective and recent advances based on “click chemistry”. Cytometry A 79, 328–337. doi:10.1002/cyto.a.21048

De Brabander, M.J., Van de Velre, R.M.L., Aerts, F.E.M., Borgers, M., Janssen, P.A.J., 1976. The Effects of Methyl [5-(2-Thienylcarbonyl)-1H-benzimidazol-2-yl]carbamate, (R 17934; NSC 238159), a New Synthetic Antitumoral Drug Interfering with Microtubules, on Mammalian Cells Cultured in Vitro. Cancer Res 36, 905–916.

Eser, P., Demel, C., Maier, K.C., Schwalb, B., Pirkl, N., Martin, D.E., Cramer, P., Tresch, A., 2014. Periodic mRNA synthesis and degradation co-operate during cell cycle gene expression. Mol Syst Biol 10, 717. doi:10.1002/msb.134886

Festuccia, N., Owens, N., Papadopoulou, T., Gonzalez, I., Tachtsidi, A., Vandoermel-Pournin, S., Gallego, E., Gutierrez, N., Dubois, A., Cohen-Tannoudji, M., Navarro, P., 2019. Transcription factor activity and nucleosome organization in mitosis. Genome Res 29, 250–260. doi:10.1101/gr.243048.118

Francis, N.J., Follmer, N.E., Simon, M.D., Aghia, G., Butler, J.D., 2009. Polycomb proteins remain bound to chromatin and DNA during DNA replication in vitro. 137, 110–122. doi:10.1016/j.cell.2009.02.017

Friman, E.T., Deluz, C., Meireles-Filho, A.C., Govindan, S., Gardeux, V., Deplancke, B., Suter, D.M., 2019. Dynamic regulation of chromatin accessibility by pluripotency transcription factors across the cell cycle. eLife 8, 380. doi:10.7554/eLife.50087

Fulco, C.P., Nasser, J., Jones, T.R., Munson, G., Bergman, D.T., Subramanian, V., Grossman, S.R., Anyoha, R., Doughty, B.R., Patwardhan, T.A., Nguyen, T.H., Kane, M., Perez, E.M., Durand, N.C., Lareau, C.A., Stamenova, E.K., Aiden, E.L., Lander, E.S., Engreitz, J.M., 2019. Activity-by-contact model of enhancer-promoter regulation from thousands of CRISPR perturbations. Nat Genet 51, 1664–1669. doi:10.1038/s41588-019-0538-0

Gonzales, K.A.U., Liang, H., Lim, Y.-S., Chan, Y.-S., Yeo, J.-C., Tan, C.-P., Gao, B., Le, B., Tan, Z.-Y., Low, K.-Y., Liou, Y.-C., Bard, F., Ng, H.-H., 2015. Deterministic Restriction on Pluripotent State Dissolution by Cell-Cycle Pathways. Cell 162, 564–579. doi:10.1016/j.cell.2015.07.001

Gu, Z., Eils, R., Schlesner, M., 2016. Complex heatmaps reveal patterns and correlations in multidimensional genomic data. Bioinformatics 32, 2847–2849. doi:10.1093/bioinformatics/btw313

Gutiérrez, M.P., MacAlpine, H.K., MacAlpine, D.M., 2019. Nascent chromatin occupancy profiling reveals locus- and factor-specific chromatin maturation dynamics behind the DNA replication fork. Genome Res 29, 1123–1133. doi:10.1101/gr.243386.118

Herzog, V.A., Fasching, N., Ameres, S.L., 2020. Determining mRNA Stability by Metabolic RNA Labeling and Chemical Nucleoside Conversion. Methods Mol Biol 2062, 169–189. doi:10.1007/978-1-4939-9822-7_9

Herzog, V.A., Reichholf, B., Neumann, T., Rescheneder, P., Bhat, P., Burkard, T.R., Wlotzka, W., Haeseler, von, A., Zuber, J., Ameres, S.L., 2017. Thiol-linked alkylation of RNA to assess expression dynamics. Nat Methods 539, 113. doi:10.1038/nmeth.4435

Kaya-Okur, H.S., Wu, S.J., Codomo, C.A., Pledger, E.S., Bryson, T.D., Henikoff, J.G., Ahmad, K., Henikoff, S., 2019. CUT&Tag for efficient epigenomic profiling of small samples and single cells. Nat Comms 10, 1930. doi:10.1038/s41467-019-09982-5

Kent, W.J., Sugnet, C.W., Furey, T.S., Roskin, K.M., Pringle, T.H., Zahler, A.M., Haussler, D., 2002. The human genome browser at UCSC. Genome Res 12, 996–1006. doi:10.1101/gr.229102

Kim, H.J., Shin, J., Lee, S., Kim, T.W., Jang, H., Suh, M.Y., Kim, J.-H., Hwang, I.-Y., Hwang, D.S., Cho, E.-J., Youn, H.-D., 2018. Cyclin-dependent kinase 1 activity coordinates the chromatin associated state of Oct4 during cell cycle in embryonic stem cells. Nucleic Acids Res 46, 6544–6560. doi:10.1093/nar/gky371

Kliszczak, A.E., Rainey, M.D., Harhen, B., Boisvert, F.M., Santocanale, C., 2011. DNA mediated chromatin pull-down for the study of chromatin replication. Sci. Rep. 1, 95. doi:10.1038/srep00095

Langmead, B., Salzberg, S.L., 2012. Fast gapped-read alignment with Bowtie 2. Nat Methods 9, 357–359. doi:10.1038/nmeth.1923

Leporcq, C., Spill, Y., Balaramane, D., Toussaint, C., Weber, M., Bardet, A.F., 2020. TFmotifView: a webserver for the visualization of transcription factor motifs in genomic regions. Nucleic Acids Res 48, W208–W217. doi:10.1093/nar/gkaa252

Liu, L., Michowski, W., Inuzuka, H., Shimizu, K., Nihira, N.T., Chick, J.M., Li, N., Geng, Y., Meng, A.Y., Ordureau, A., Kołodziejczyk, A., Ligon, K.L., Bronson, R.T., Polyak, K., Harper, J.W., Gygi, S.P., Wei, W., Sicinski, P., 2017. G1 cyclins link proliferation, pluripotency and differentiation of embryonic stem cells. Nat. Cell Biol. 19, 177–188. doi:10.1038/ncb3474

Liu, L., Michowski, W., Kolodziejczyk, A., Sicinski, P., 2019. The cell cycle in stem cell proliferation, pluripotency and differentiation. Nat. Cell Biol. 21, 1060–1067. doi:10.1038/s41556-019-0384-4

Liu, Y., Pelham-Webb, B., Di Giammartino, D.C., Li, J., Kim, D., Kita, K., Saiz, N., Garg, V., Doane, A., Giannakakou, P., Hadjantonakis, A.-K., Elemento, O., Apostolou, E., 2017. Widespread Mitotic Bookmarking by Histone Marks and Transcription Factors in Pluripotent Stem Cells. Cell Rep 19, 1283–1293. doi:10.1016/j.celrep.2017.04.067

Lizio, M., Abugessaisa, I., Noguchi, S., Kondo, A., Hasegawa, A., Hon, C.C., de Hoon, M., Severin, J., Oki, S., Hayashizaki, Y., Carninci, P., Kasukawa, T., Kawaji, H., 2019. Update of the FANTOM web resource: expansion to provide additional transcriptome atlases. Nucleic Acids Res 47, D752–D758. doi:10.1093/nar/gky1099

Love, M.I., Huber, W., Anders, S., 2014. Moderated estimation of fold change and dispersion for RNA-seq data with DESeq2. Genome Biol 15, 550. doi:10.1186/s13059-014-0550-8

Ly, T., Endo, A., Lamond, A.I., 2015. Proteomic analysis of the response to cell cycle arrests in human myeloid leukemia cells. eLife 4, M111.013680. doi:10.7554/eLife.04534

Ly, T., Whigham, A., Clarke, R., Brenes-Murillo, A.J., Estes, B., Madhessian, D., Lundberg, E., Wadsworth, P., Lamond, A.I., 2017. Proteomic analysis of cell cycle progression in asynchronous cultures, including mitotic subphases, using PRIMMUS. eLife 6. doi:10.7554/eLife.27574

Macheret, M., Halazonetis, T.D., 2019. Monitoring early S-phase origin firing and replication fork movement by sequencing nascent DNA from synchronized cells. Nat Protoc 14, 51–67. doi:10.1038/s41596-018-0081-y

Martin, M., 2011. Cutadapt removes adapter sequences from high-throughput sequencing reads. EMBnet.journal 17, 10–12.

Michowski, W., Chick, J.M., Chu, C., Kolodziejczyk, A., Wang, Y., Suski, J.M., Abraham, B., Anders, L., Day, D., Dunkl, L.M., Li Cheong Man, M., Zhang, T., Laphanuwat, P., Bacon, N.A., Liu, L., Fassl, A., Sharma, S., Otto, T., Jecrois, E., Han, R., Sweeney, K.E., Marro, S., Wernig, M., Geng, Y., Moses, A., Li, C., Gygi, S.P., Young, R.A., Sicinski, P., 2020. Cdk1 Controls Global Epigenetic Landscape in Embryonic Stem Cells. Mol Cell 78, 459–476.e13. doi:10.1016/j.molcel.2020.03.010

Murray, A.W., Kirschner, M.W., 1989. Dominoes and clocks: the union of two views of the cell cycle. Science 246, 614–621. doi:10.1126/science.2683077

Neumann, T., Herzog, V.A., Muhar, M., Haeseler, von, A., Zuber, J., Ameres, S.L., Rescheneder, P., 2019. Quantification of experimentally induced nucleotide conversions in high-throughput sequencing datasets. BMC Bioinformatics 20, 258. doi:10.1186/s12859-019-2849-7

Otto, T., Sicinski, P., 2017. Cell cycle proteins as promising targets in cancer therapy. Nat. Rev. Cancer 17, 93–115. doi:10.1038/nrc.2016.138

Owens, N., Papadopoulou, T., Festuccia, N., Tachtsidi, A., Gonzalez, I., Dubois, A., Vandormael-Pournin, S., Nora, E.P., Bruneau, B.G., Cohen-Tannoudji, M., Navarro, P., 2019. CTCF confers local nucleosome resiliency after DNA replication and during mitosis. eLife 8, 281. doi:10.7554/eLife.47898

Pauklin, S., Vallier, L., 2013. The cell-cycle state of stem cells determines cell fate propensity. 155, 135–147. doi:10.1016/j.cell.2013.08.031

Petryk, N., Dalby, M., Wenger, A., Stromme, C.B., Strandsby, A., Andersson, R., Groth, A., 2018. MCM2 promotes symmetric inheritance of modified histones during DNA replication. Science 361, 1389–1392. doi:10.1126/science.aau0294

Pintacuda, G., Wei, G., Roustan, C., Kirmizitas, B.A., Solcan, N., Cerase, A., Castello, A., Mohammed, S., Moindrot, B., Nesterova, T.B., Brockdorff, N., 2017. hnRNPK Recruits PCGF3/5-PRC1 to the Xist RNA B-Repeat to Establish Polycomb-Mediated Chromosomal Silencing. Mol Cell 68, 955–969.e10. doi:10.1016/j.molcel.2017.11.013

Pontarin, G., Gallinaro, L., Ferraro, P., Reichard, P., Bianchi, V., 2003. Origins of mitochondrial thymidine triphosphate: dynamic relations to cytosolic pools. Proc Natl Acad Sci USA 100, 12159–12164. doi:10.1073/pnas.1635259100

Ramírez, F., Ryan, D.P., Grüning, B., Bhardwaj, V., Kilpert, F., Richter, A.S., Heyne, S., Dündar, F., Manke, T., 2016. deepTools2: a next generation web server for deep-sequencing data analysis. Nucleic Acids Res 44, W160–5. doi:10.1093/nar/gkw257

Reverón-Gómez, N., González-Aguilera, C., Stewart-Morgan, K.R., Petryk, N., Flury, V., Graziano, S., Johansen, J.V., Jakobsen, J.S., Alabert, C., Groth, A., 2018. Accurate Recycling of Parental Histones Reproduces the Histone Modification Landscape during DNA Replication. Mol Cell 72, 239–249.e5. doi:10.1016/j.molcel.2018.08.010

Sakaue-Sawano, A., Kurokawa, H., Morimura, T., Hanyu, A., Hama, H., Osawa, H., Kashiwagi, S., Fukami, K., Miyata, T., Miyoshi, H., Imamura, T., Ogawa, M., Masai, H., Miyawaki, A., 2008. Visualizing spatiotemporal dynamics of multicellular cell-cycle progression. 132, 487–498. doi:10.1016/j.cell.2007.12.033

Singh, A.M., Chappell, J., Trost, R., Lin, L., Wang, T., Tang, J., Matlock, B.K., Weller, K.P., Wu, H., Zhao, S., Jin, P., Dalton, S., 2013. Cell-cycle control of developmentally regulated transcription factors accounts for heterogeneity in human pluripotent cells. Stem Cell Reports 1, 532–544. doi:10.1016/j.stemcr.2013.10.009

Singh, A.M., Sun, Y., Li, L., Zhang, W., Wu, T., Zhao, S., Qin, Z., Dalton, S., 2015. Cell-Cycle Control of Bivalent Epigenetic Domains Regulates the Exit from Pluripotency. Stem Cell Reports 5, 323–336. doi:10.1016/j.stemcr.2015.07.005

Skinner, S.O., Xu, H., Nagarkar-Jaiswal, S., Freire, P.R., Zwaka, T.P., Golding, I., 2016. Single-cell analysis of transcription kinetics across the cell cycle. eLife 5, e59928. doi:10.7554/eLife.12175

Stadler, M.B., Murr, R., Burger, L., Ivanek, R., Lienert, F., Schöler, A., van Nimwegen, E., Wirbelauer, C., Oakeley, E.J., Gaidatzis, D., Tiwari, V.K., Schübeler, D., 2011. DNA-binding factors shape the mouse methylome at distal regulatory regions. Nature 480, 490–495. doi:10.1038/nature10716

Stewart-Morgan, K.R., Reverón-Gómez, N., Groth, A., 2019. Transcription Restart Establishes Chromatin Accessibility after DNA Replication. Mol Cell 75, 284–297.e6. doi:10.1016/j.molcel.2019.04.033

Teves, S.S., An, L., Hansen, A.S., Xie, L., Darzacq, X., Tjian, R., 2016. A dynamic mode of mitotic bookmarking by transcription factors. eLife 5. doi:10.7554/eLife.22280

Trcek, T., Larson, D.R., Moldón, A., Query, C.C., Singer, R.H., 2011. Single-molecule mRNA decay measurements reveal promoter- regulated mRNA stability in yeast. 147, 1484–1497. doi:10.1016/j.cell.2011.11.051

Vassilev, L.T., Tovar, C., Chen, S., Knezevic, D., Zhao, X., Sun, H., Heimbrook, D.C., Chen, L., 2006. Selective small-molecule inhibitor reveals critical mitotic functions of human CDK1. Proc Natl Acad Sci USA 103, 10660–10665. doi:10.1073/pnas.0600447103

Voichek, Y., Bar-Ziv, R., Barkai, N., 2016. Expression homeostasis during DNA replication. Science 351, 1087–1090. doi:10.1126/science.aad1162

Yan, J., Chen, S.-A.A., Local, A., Liu, T., Qiu, Y., Dorighi, K.M., Preissl, S., Rivera, C.M., Wang, C., Ye, Z., Ge, K., Hu, M., Wysocka, J., Ren, B., 2018. Histone H3 lysine 4 monomethylation modulates long-range chromatin interactions at enhancers. Cell Res 28, 204–220. doi:10.1038/cr.2018.1

Yiangou, L., Grandy, R.A., Morell, C.M., Tomaz, R.A., Osnato, A., Kadiwala, J., Muraro, D., Garcia-Bernardo, J., Nakanoh, S., Bernard, W.G., Ortmann, D., McCarthy, D.J., Simonic, I., Sinha, S., Vallier, L., 2019. Method to Synchronize Cell Cycle of Human Pluripotent Stem Cells without Affecting Their Fundamental Characteristics. Stem Cell Reports 12, 165–179. doi:10.1016/j.stemcr.2018.11.020

Zhang, H., Emerson, D.J., Gilgenast, T.G., Titus, K.R., Lan, Y., Huang, P., Zhang, D., Wang, H., Keller, C.A., Giardine, B., Hardison, R.C., Phillips-Cremins, J.E., Blobel, G.A., 2019. Chromatin structure dynamics during the mitosis-to-G1 phase transition. Nature 576, 158–162. doi:10.1038/s41586-019-1778-y

Zhang, Y., Liu, T., Meyer, C.A., Eeckhoute, J., Johnson, D.S., Bernstein, B.E., Nusbaum, C., Myers, R.M., Brown, M., Li, W., Liu, X.S., 2008. Model-based analysis of ChIP-Seq (MACS). Genome Biol 9, R137. doi:10.1186/gb-2008-9-9-r137

