## Supplemental files for "An integrated approach to study chromatin and transcriptional dynamics throughout the cell cycle without pharmacological inhibition"

### Supplemental figures
### Shlyueva et al.

Supplemental Figure 1
Supplemental Figure 2
Supplemental Figure 3
Supplemental Figure 4
Supplemental Figure 5

Supplemental Figure S1

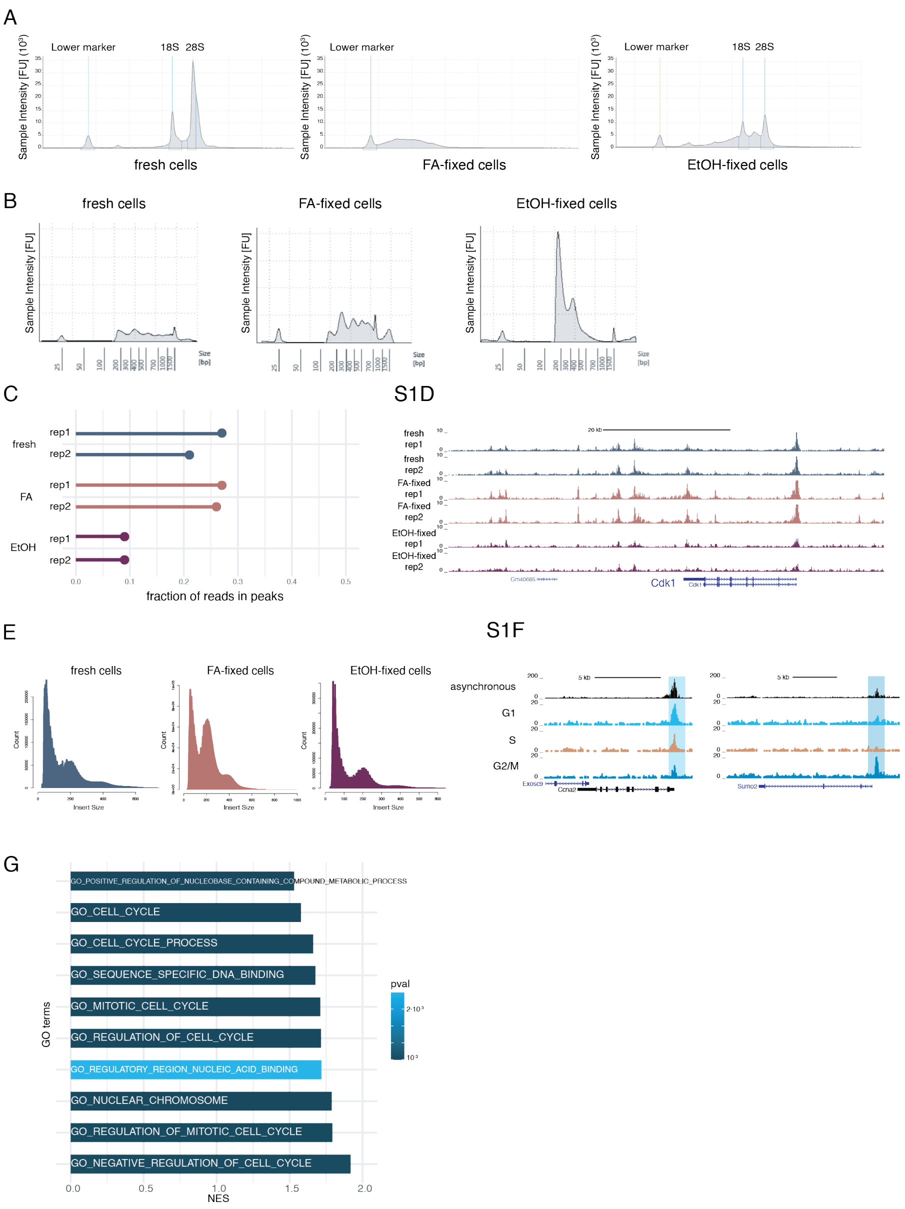


**Supplemental Figure S1. Integration of fixing and cell cycle staining with profiling of chromatin accessibility and gene expression.** (A) Intensities of the major ribosomal RNA (rRNA) bands and degradation products after different types of fixation. The leftmost peak is the lower marker. (B) Size distribution (bp) of ATAC-seq libraries from fresh, FA- and EtOH-fixed cells. (C) Lollipop plot showing the ratio of reads in ATAC-seq peak to all reads in each sample. (D) UCSC genome browser screenshot of ATAC-seq read densities in fresh, FA- and EtOH-fixed mESC. (E) Insert size distribution of ATAC-seq reads in libraries from fresh, FA- and EtOH-fixed mESC. (F) UCSC genome browser screenshots of asynchronous, G1, S and G2/M ATAC-seq accessibility tracks. G1-specific peak at *Ccna2* promoter highlighted in light blue (left) and G2/M-specific peak at *Sumo2* – in dark blue (right). (G) Bar plots showing the normalized enrichment scores and p-values for terms in gene ontology sets (C5) from MSigDB for identified cycling genes. The top-10 terms based on p-value are shown.

Supplemental Figure S2

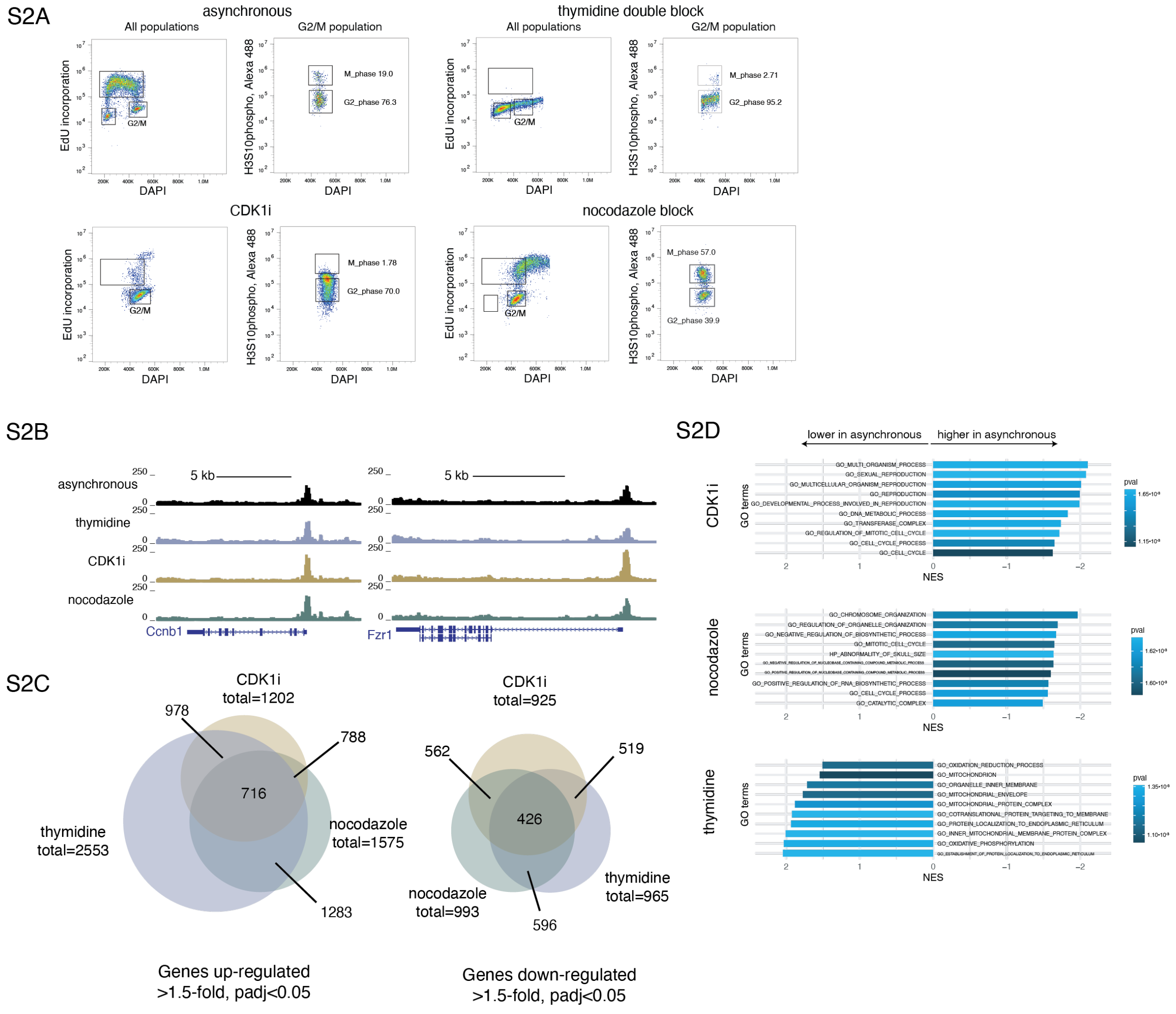


**Supplemental Figure S2. Chromatin accessibility and gene expression in drug synchronized mESC.** (A) Representative flow cytometry profiles of mESC: asynchronous or synchronized with double thymidine block, CDK1 inhibitor or nocodazole. G2 and M phases were separated by co-staining for H3S10phosho. The indicated percentages show the proportion of G2/M population. The FACS profile gates for nocodazole-treated cells were adjusted to include two distinct populations. (B) UCSC genome browser screenshots of ATAC-seq tracks from asynchronous, thymidine-, CDKi- or nocodazole-treated cells. Peaks at *Ccnb1* and *Fzr1* promoters are shown. (C)Venn diagrams showing the overlap of up-regulated (left) or down-regulated (right) genes after drug treatments. (D) Bar plots showing the normalized enrichment scores and p-values for terms in gene ontology sets (C5) from MSigDB for identified cycling genes. The top-10 terms based on p-value are shown.

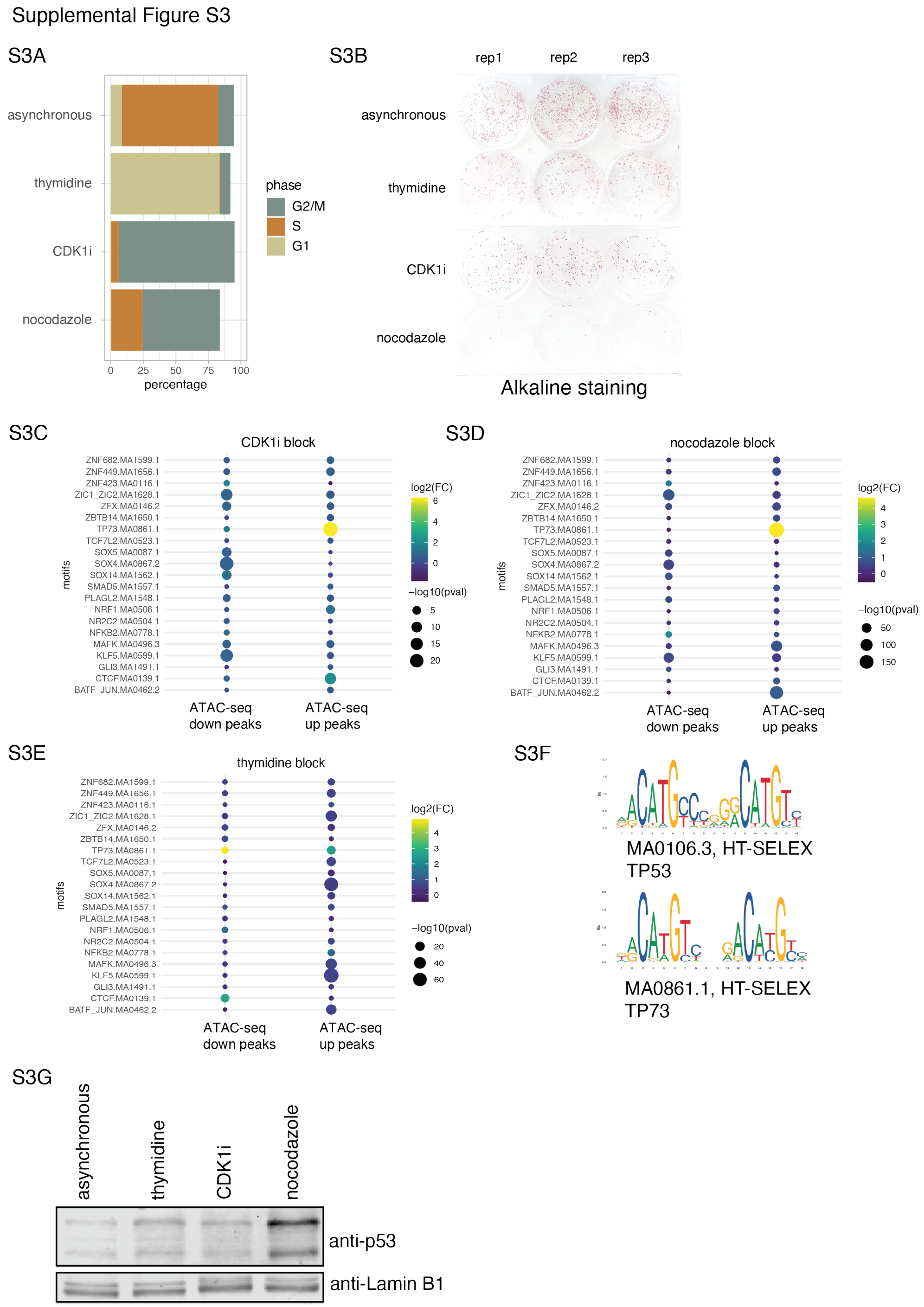


**Supplemental Figure S3. Effect of synchronization on mESCs colony-forming ability and motif analysis for open chromatin regions.** (A) Cell cycle phase distribution for asynchronous cells and synchronized cells before the colony-forming assay and Western blot. (B) The colony-forming assay for mESC cells with and without inhibitors in three technical replicates. (C) (D) (E) Heatmap presenting enrichment (color-coded log2 fold-changes) and significance (-log10(p-value) proportional to sizes of circles) for transcription factors' motifs compared to random genomic regions. (F) The sequence logo for p53 and p73 motifs enriched in ATAC peaks after synchronization. (G) Western blot shows abundance of p53 in mESC with and without synchronization in comparison to Lamin B1.

Supplemental Figure S4

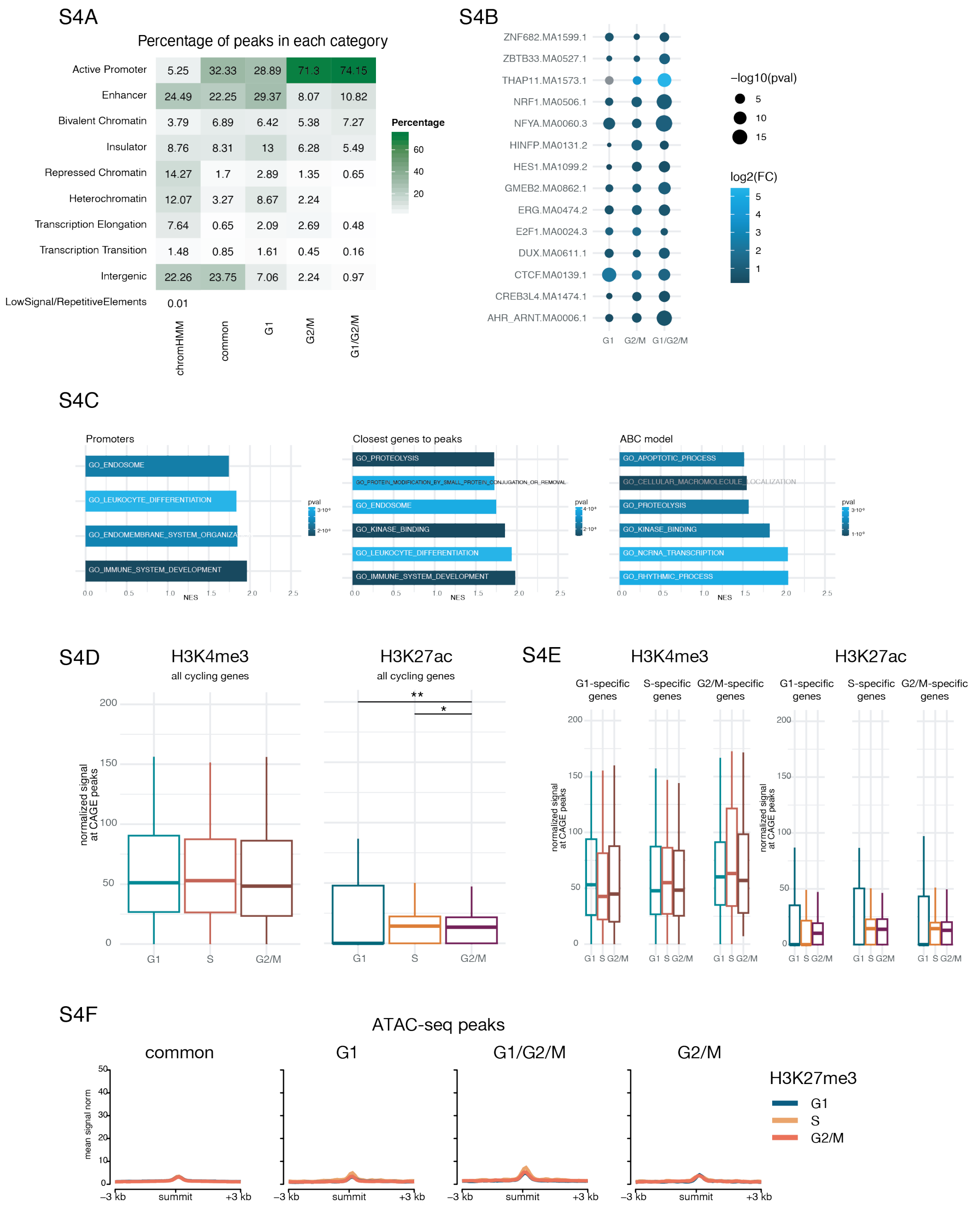


**Supplemental Figure S4. Chromatin states of cell cycle-dependent accessible regions and genes.** (A) Heatmap showing percentage of co-occurrences for common or cell cycle-specific ATAC peaks with chromatin states (chromHMM) [45] as well as genome-wide frequency of chromatin states. (B) Heatmap presenting enrichment (color-coded log2 fold-changes) and significance (-log10(p-value) proportional to sizes of circles) for transcription factors' motifs compared to random genomic regions. (C) Bar plots showing the normalized enrichment scores and p-values for terms in gene ontology sets (C5) from MSigDB for genes with cell cycle-dependent open region at their promoters (left), closest to them (middle) or assigned to peaks via ABC model (right). The significant terms based on p-value are shown. (D) Box plots depicting the distribution of normalized reads counts for H3K4me3 and H3K27ac over promoters of cycling genes (defined based on CAGE data). Wilcoxon rank-sum test: *p-value<0.05, **p-value<0.01. The box designates the 25th and 75th percentiles and is divided by the median; the whiskers extend to 5th and 95th percentiles. (E) Similar to (D), but the cycling genes are broken down into cell cycle-specific categories. (F) Average profiles over different categories of ATAC-seq peaks (± 3 kb) for H3K27me3 in G1, S and G2/M cells sorted from unperturbed mESC.

Supplemental Figure S5

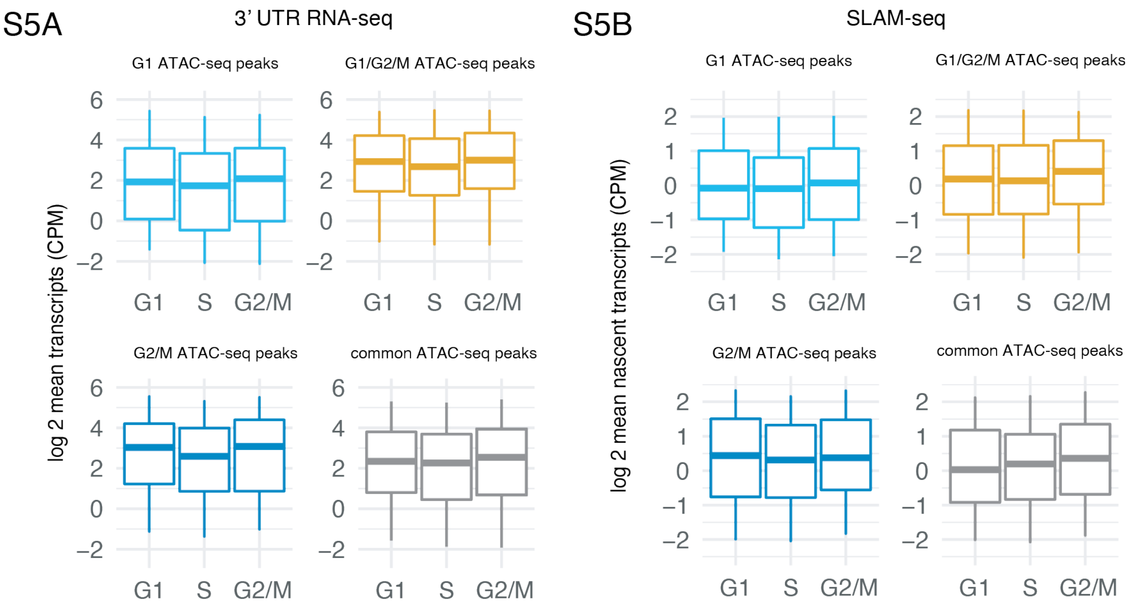


**Supplemental Figure S5. Correlation between cell cycle-dependent ATAC-seq and RNA-seq or SLAM-seq.** (A) Box plots showing counts per million reads mapped (CPM, log2 scale) for steady transcripts' levels or (B) nascent transcripts of genes closest to cell cycle-specific ATAC-seq peaks. The box designates the 25$^{th}$ and 75$^{th}$ percentiles and is divided by the median; the whiskers extend to 5$^{th}$ and 95$^{th}$ percentiles.
